# Personalizing a computational upper body model improves kinematic tracking in high range-of-motion arm movements

**DOI:** 10.1101/2024.06.12.598739

**Authors:** Jennifer N. Maier, Nicholas A. Bianco, Carmichael F. Ong, Julie Muccini, Ellen Kuhl, Scott L. Delp

## Abstract

Musculoskeletal models of the shoulder are needed to understand the mechanics of overhead motions. Existing models can be scaled to represent the size of an individual person, but the kinematics are generic. We introduce a method to personalize the shoulder kinematics of a computational model of the upper body that defines the orientations of the clavicle and scapula based on glenohumeral joint angles. During five static calibration poses, we palpate and measure the orientation of the scapula. We explore the importance of representing shoulder elevation by introducing clavicle elevation as a degree of freedom that is independent of the glenohumeral angles. For ten subjects, we record the five calibration poses, ten additional static poses, and dynamic arm raises using optical motion capture. We examine the data using a dynamically-constrained inverse kinematics analysis. Personalization, independent clavicle elevation, and both in combination reduce the average upper body marker tracking error compared to the generic model in the static poses (26 mm to 17-20 mm) and in the dynamic trials (22 mm to 14-17 mm). Only personalization reduces the average scapula marker error (51 mm to 36-38 mm) and scapula axis-angle error (15° to 10°) in the static poses, and in the dynamic trials at instances that best match the static poses (53 mm to 37-40 mm, 15° to 9°). Our results show that personalizing upper body models improves kinematic tracking. We provide our experimental data, model, and methods to allow researchers to reproduce and build upon our results.

## 1. Introduction

Realistic computational models of shoulder kinematics are essential when analyzing the biomechanics of the upper body during high range-of-motion arm movements. Understanding these types of movements is relevant in a variety of applications, such as analyzing rehabilitation after shoulder injuries and understanding overhead athletes like volleyball hitters or baseball pitchers, who perform high range-of-motion movements (Baena-Raya et al., 2021; Hashimoto et al., 2023).

A challenge for modeling shoulder kinematics is the complexity of the shoulder. It consists of four joints: the sternoclavicular joint, acromioclavicular joint, glenohumeral joint, and scapulothoracic articulation. These joints form a closed loop, which makes modeling and understanding the dynamics of the shoulder difficult. Biomechanical analysis is further complicated by difficulties in obtaining accurate experimental measurements of shoulder motions as much of the movement takes place under a layer of soft tissues. Methods like bone pins or fluoroscopy used to obtain ground truth measurements of scapular motion are infeasible for routine screenings and analyses of athletic motions (Giphart et al., 2013; Ivester et al., 2015; Ludewig et al., 2009; Nicholson et al., 2017a; Seth et al., 2016; Yanagawa et al., 2008). Methods to improve the accuracy of measuring and modeling shoulder kinematics from optical motion measurements would thus be valuable.

Several approaches have been used to model the shoulder. Van der Helm (1994) models the scapulothoracic gliding plane in a finite element model used for kinetics analysis. This approach was later implemented in a musculoskeletal model that traced a point on the scapula that slides on an ellipsoid surface representing the thoracic cage (Seth et al., 2016). This modeling approach enables flexible positioning of the scapula during a variety of motions but requires measurement of the position of the scapula to achieve realistic motion tracking.

An alternative approach for modeling shoulder motions is to implement the shoulder rhythm, which describes the position and orientation of the scapula and clavicle based on the glenohumeral joint angles. A variety of valuable biomechanical models have implemented the shoulder rhythm (De Groot and Brand, 2001; Grewal and Dickerson, 2013; Holzbaur et al., 2005; Makhsous, 1999; Saul et al., 2015; Xu et al., 2014), yet, a comparison of these models shows that none performs consistently better than others in representing static pose orientation measurements (Xu et al., 2016). Personalizing the shoulder rhythm dependencies based on calibration measurements can improve the tracking of scapular orientation, as shown by the agreement with ground truth fluoroscopic measurements (Nicholson et al., 2017b).

To further develop an approach to personalize a model of the shoulder, we build on the upper body model that was first described by Holzbaur et al. (2005) and later updated by Saul et al. (2015), which implements the shoulder rhythm by coupling the sternoclavicular and acromioclavicular joint angles to the glenohumeral angles using linear relationships. This widely used model derives these linear relationships using regression on data from ten healthy young participants (De Groot and Brand, 2001). Personalizing the shoulder rhythm may improve accuracy in analyzing an individual, and in particular in overhead athletes who display substantial differences in resting position and in scapulohumeral rhythm between their dominant and non-dominant shoulders (Pascoal et al., 2023; Ribeiro and Pascoal, 2013). A limitation of shoulder rhythm models is that they fail to accurately replicate elevation and depression of the shoulders. These motions are key parts of the movement patterns of many overhead athletes, and capturing these features is critical for accurate analysis of these athletes.

The objective of this work is to evaluate whether personalizing the shoulder constraints in a computational upper body model (Holzbaur et al., 2005; Saul et al., 2015) based on five calibration poses can reduce marker tracking errors in the kinematic analysis of shoulder motions. We also investigate the effect of allowing independent clavicle elevation instead of constraining it within the shoulder rhythm. We focus on scapular kinematics because they are crucial for accurate shoulder motion analysis.

## 2. Methods

We collected experimental data and used them to modify an upper body model by personalizing the shoulder rhythm and allowing for independent clavicle elevation motion. We computed kinematics in a direct collocation framework and evaluated the effects of the model modifications compared to the generic model.

### 2.1. Experimental Data

We recorded data in a laboratory setting with a 16-camera optical motion capture system at 200 Hz (Motion Analysis Corporation, Santa Rosa, CA, USA) measuring retroreflective markers placed on the upper body (Fig. 1). We recorded motions in 10 healthy young individuals, eight of whom were right-handed, and two were left-handed. Half of the subjects were female (25.8 ± 2.0 years, 1.68 ± 0.05 m, 64.0 ± 7.0 kg), the other half were male (31.8 ± 2.2 years, 1.86 ± 0.09 m, 79.8 ± 7.5 kg). For each participant, we recorded five static calibration trials for each shoulder, with different glenohumeral joint angles, including the anatomical position, maximal abduction, maximal flexion, maximal horizontal abduction, and maximal horizontal adduction (Fig. 2A). We palpated the positions of the trigonum spinae, angulus inferior, and angulus acromialis of the scapula, placed an L-shaped scapula locator on the identified locations, and held it in place during a one-second long recording (Fig. 2B).

**Figure 1.**
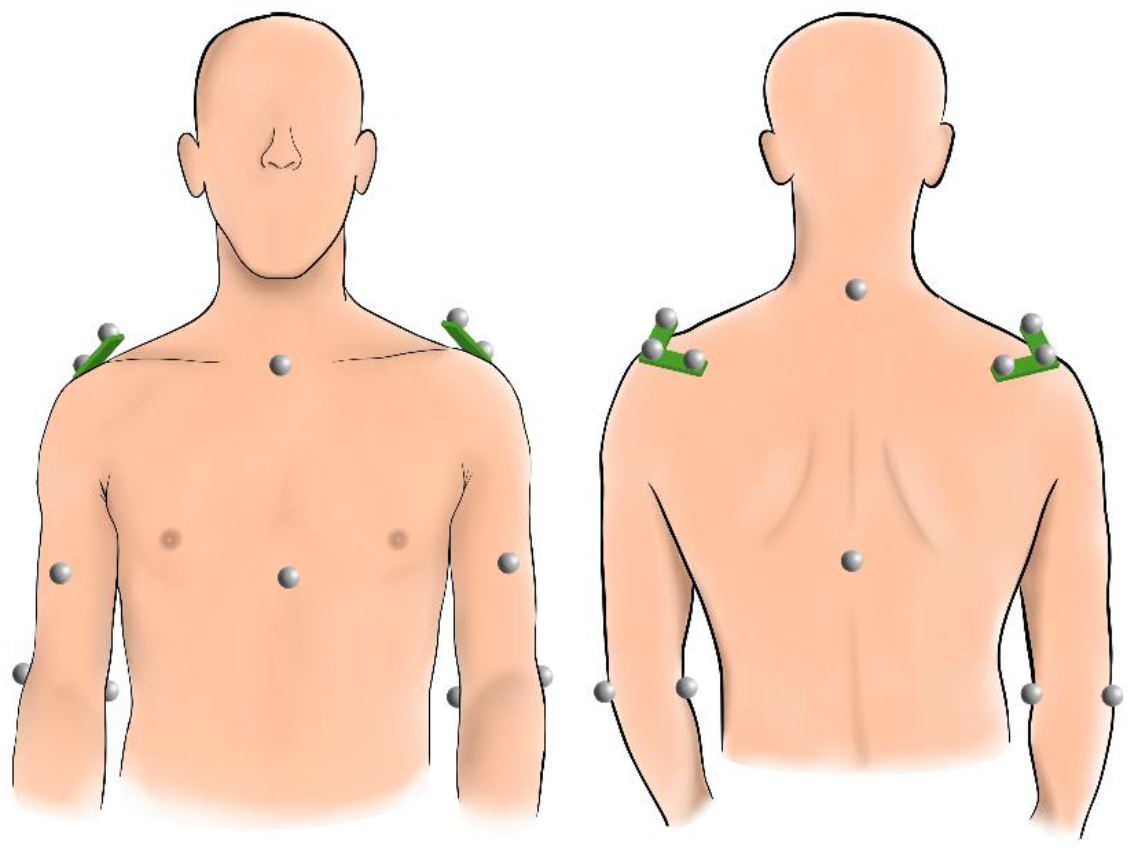
Marker setup with markers on the torso (7th cervical vertebra, the 8th thoracic vertebra, the suprasternal notch, and the xiphoid process), on the arms (lateral and medial epicondyle of the humerus), and on the shoulders (three-marker acromion cluster with the central marker located on the flat part of the acromion, one arm of the cluster along the spine of the scapula, and the other arm of the cluster pointing forwards into the air to allow the cluster to move with shoulder motion).

**Figure 2.**
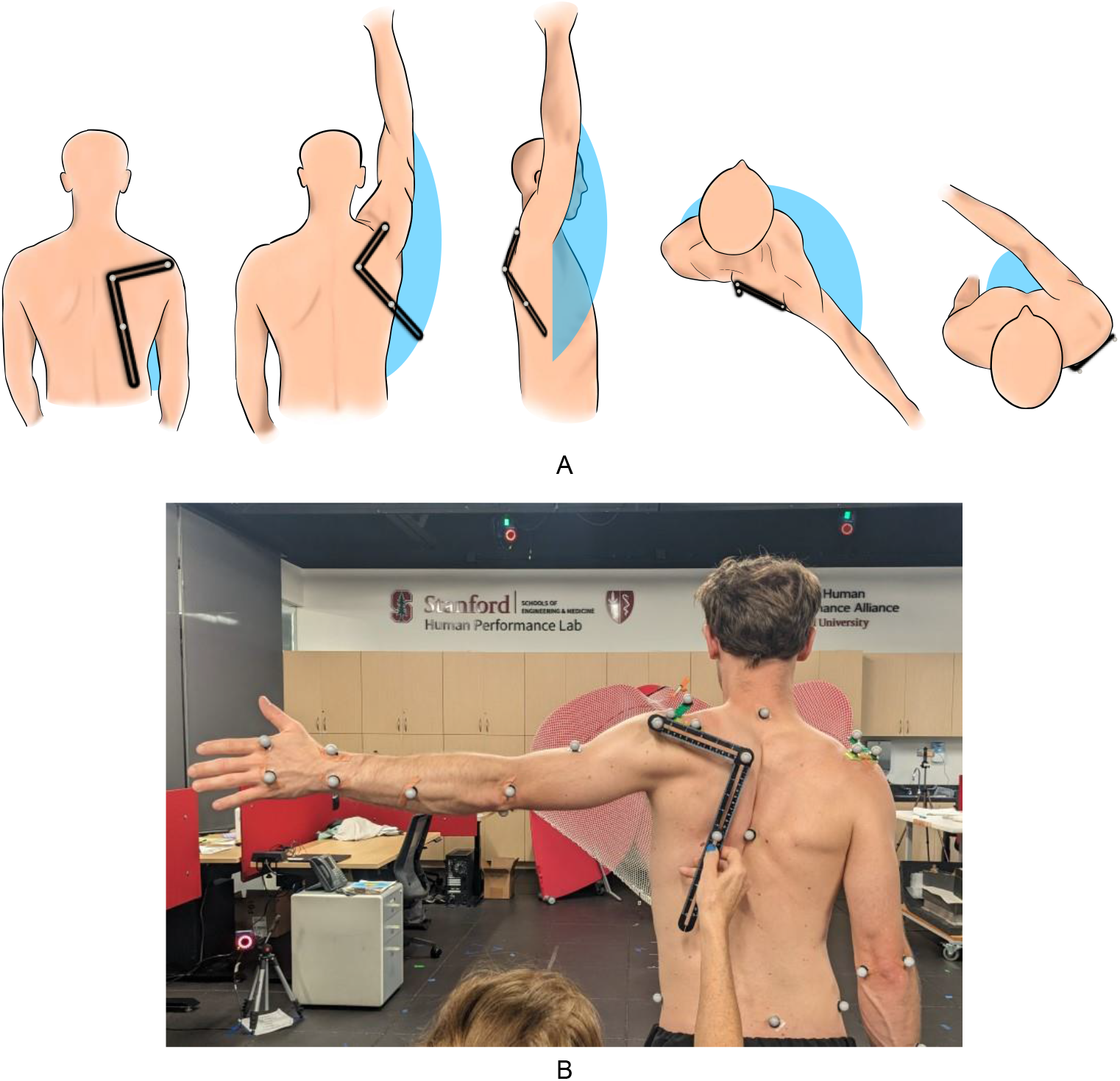
Calibration. A) Five calibration poses for the right arm: the anatomical position, maximal abduction, maximal flexion, maximal horizontal abduction, and maximal horizontal adduction, from left to right. The black L-shape represents the scapula locator, and the blue areas visualize the humerus elevation angle in the respective body plane. B) Measurement of the scapula position at maximal horizontal abduction with the L-shaped scapula locator shown in black. The occupational therapist places the locator on three palpated points on the scapula and kneels behind the participant for recording to minimize marker occlusion.

To test the effects of model personalization, we then recorded a variety of static poses during which we palpated and measured the scapula of the dominant arm. The measured poses include shoulder abduction at 45°, 90°, and 135°; maximal shoulder extension; shoulder flexion at 45°, 90°, and 135°; horizontal abduction of 45° at 90° flexion; and maximal elevation and depression of the shoulders. After the static trials, the participants performed three trials each of dynamic shoulder abduction, flexion, and horizontal abduction at 90° shoulder flexion. For one subject, we did not record static elevation and depression, and for another subject, one dynamic horizontal abduction trial had missing data.

### 2.2. Musculoskeletal Model

We personalized the upper body biomechanical model by Saul et al. (2015). In this generic model, the rotations of the glenohumeral joint were defined in the globe coordinate system as described in Doorenbosch et al. (2003), including the elevation plane angle, elevation angle, and rotation angle. Based on the findings of De Groot and Brand (2001), the generic model implemented kinematic constraints defining the motion of the clavicle protraction and elevation, and the scapula internal rotation, upward rotation, and posterior tilting; each constraint was a linear function of the humeral elevation angle and neglected additional dependencies of these joint angles on the glenohumeral elevation plane angle found in De Groot and Brand (2001). We included these additional dependencies, such that the generic model described each clavicle and scapula coordinate by a bivariate constraint dependent on the elevation plane angle and the glenohumeral elevation plane angle.

### 2.3. Model Modifications

We investigated two changes to the shoulder joint model: first, personalizing the linear constraints based on the calibration measurements, and second, removing the clavicle elevation constraint and including clavicle elevation as an additional degree-of-freedom. We thus compared four models: the fully constrained generic model with the bivariate constraints as described in section 2.2, the fully constrained model with personalized constraints, the model with generic bivariate constraints and independent clavicle elevation, and the model with both personalized constraints and independent clavicle elevation.

### 2.4. Model Personalization

We updated the shoulder joint coordinate systems to follow the recommendations of the International Society of Biomechanics (Wu et al., 2005; Xu et al., 2012). Table A.1 in Appendix A shows the differences between the coordinate systems.

Using the OpenSim Scale Tool, we scaled the model to each subject’s anthropometry using the marker coordinates from the static measurement in the anatomical pose (Delp et al., 2007; Seth et al., 2018). Since scaling with only the experimental marker data in a single pose often resulted in excessive humerus lengths, we updated this scaling factor by estimating the glenohumeral rotation center in all five calibration poses using the SCoRE method and calculating the mean scaling factor (Monnet et al., 2007). Because some of the bones were scaled anisotropically, we updated the joint coordinate frames after scaling.

The dependencies of the clavicle and scapula orientations on the humerus orientation were implemented using OpenSim’s *CoordinateCouplerConstraint*. For the sternoclavicular joint angles and acromioclavicular joint angles, a linear bivariate function of the form

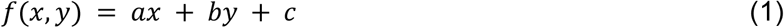

described the angle with respect to the glenohumeral elevation plane angle *x* and glenohumeral elevation angle *y*.

To personalize the model to each participant, we calculated the clavicle, scapula, and humerus orientations relative to the thorax in all five calibration poses. Note that this computation was performed on the marker data directly, only using the origins and orientations of the model’s joint coordinate systems. For each dependent sternoclavicular and acromioclavicular joint angle, we then fit a bilinear plane to the computed orientations using least squares. We updated the parameters *a, b*, and *c* of the bivariate *CoordinateCouplerConstraints* (Eq. 1) with the slopes and intercept of each plane to create the personalized model.

### 2.5. Independent clavicle elevation motion

A limitation of implementing the shoulder rhythm was that the model could not replicate shoulder motions that were independent of humeral elevation like elevating the shoulders (Seth et al., 2016). Thus, we investigated the feasibility of introducing an independent coordinate for clavicle elevation while keeping all other shoulder joint constraints in place.

Since our marker-based recordings only contained a single marker on the sternal notch and a marker cluster on the flat part of the acromion, we did not have sufficient information to track clavicle elevation. We thus estimated the scapula position from an acromion marker cluster. This method estimated the scapula position and orientation relative to the acromion marker cluster in the three body planes based on the orientation of the humerus and formed a weighted average of these three estimates. It then computed the global estimate for the scapula position and orientation by registering to the current measurement of the acromion marker cluster. Using this scapula estimate, we computed the position of the acromioclavicular joint that marks the distal end of the clavicle and used this position as a virtual marker to track the clavicle elevation during the inverse kinematics computation.

### 2.6. Simulations

We added actuators to all joints of the models and used OpenSim Moco to generate simulations that minimized the error between experimental and model marker positions and the squared actuator controls using direct collocation optimal control (Dembia et al., 2020). The torso markers were weighted highest to prevent excessive lumbar bending, which may have occurred if the shoulder joint definition could not achieve good tracking of both arm and torso markers. When tracking the clavicle elevation, we weighted the virtual marker at the acromioclavicular joint higher than the markers on the arms to achieve smoother shoulder elevation changes. We solved the tracking problems with CasADi (Andersson et al., 2019) and IPOPT (Wächter and Biegler, 2006) using a Hermite-Simpson transcription scheme (Betts, 2010). Simulation details are given in Appendix B.

### 2.7. Validation Approach

We validated our kinematics analysis following common guidelines (Hicks et al., 2015). In all static trials where a palpated ground truth measurement of the scapula position was available, we computed the root mean squared (RMS) errors between experimental and model marker trajectories. We did this first for all markers, and then only for the three scapula markers. Furthermore, we calculated the axis-angle error between the palpated and estimated scapula orientation (Al Borno et al., 2022).

In the dynamic trials, where no palpated scapula information was available, we computed the RMS errors between experimental and model marker trajectories for all anatomical markers measured during those trials. To obtain an estimate of how well the shoulder models perform regarding clavicle and scapula orientation during each dynamic movement, we compared the scapula orientation at time points of the dynamic trials at which the marker data best match the static trials. To find the best matching time points in the dynamic trials, we first used the four torso markers to rigidly register the marker data of each dynamic trial to each static trial. We then found the frame at which the two elbow markers of the moving arm of the dynamic trial had the smallest Euclidean distance to the elbow markers of the static trial.

For the dynamic abduction trials, we compared with the static poses at 45°, 90°, and 135° abduction; for the dynamic flexion trials, we compared with the static measurements at maximal extension, 45°, 90°, and 135° flexion; and for the dynamic horizontal abduction trials, we compared with the trials at 90° flexion, horizontal abduction of 45° at 90° flexion, and 90° abduction. At each best matching time point, we computed the average scapula marker error and the axis-angle error between the palpated static measurements and the estimated scapula orientation of the dynamic trial.

### 2.8. Statistical Testing

We used the Python libraries statsmodels and SciPy stats for all statistical computations (Seabold and Perktold, 2010; Virtanen et al., 2020). Shapiro-Wilk tests did not show evidence of non-normality in any of our comparisons (BenSaïda, 2014). We fit linear mixed models to test the effects of the changes we made to the generic model while controlling for the within-subject nature of the four compared models. Our dependent variable was the marker RMS error, and our fixed terms effects were personalizing the constraint equations, using an independent coordinate for clavicle elevation, and the interaction between these model modifications. We considered *p-*values < 0.05 significant. In cases where an interaction effect was detected, we further investigated individual between-model changes with dependent-samples *t*-tests controlling for the false discovery rate (Benjamini and Hochberg, 1995).

## 3. Results

Personalizing the model and allowing independent clavicle elevation motion improved the upper body kinematic tracking during static trials, but only personalization led to a better tracking of scapular kinematics. To visualize the improvements, we compared the simulation results with a “target model”, which had an unconstrained shoulder and tracked the palpated scapula markers in the kinematics computation to determine the best possible clavicle and scapula angles (Fig. 3). In high range-of-motion poses (first two rows), all modifications led to an arm position that was closer to that of the target model when compared to the generic model, but only the models with personalized constraints improved the scapula fit with the target model. The fully-constrained generic model matched better with the target model for shoulder elevation (third row), and the fully-constrained personalized model matched better for depression (fourth row), but neither was able to track both of these poses well. Both models with independent clavicle elevation motion achieved a better fit to the shoulder height in elevation and depression, but the scapula orientation of the personalized model was closer to the target model.

**Figure 3.**
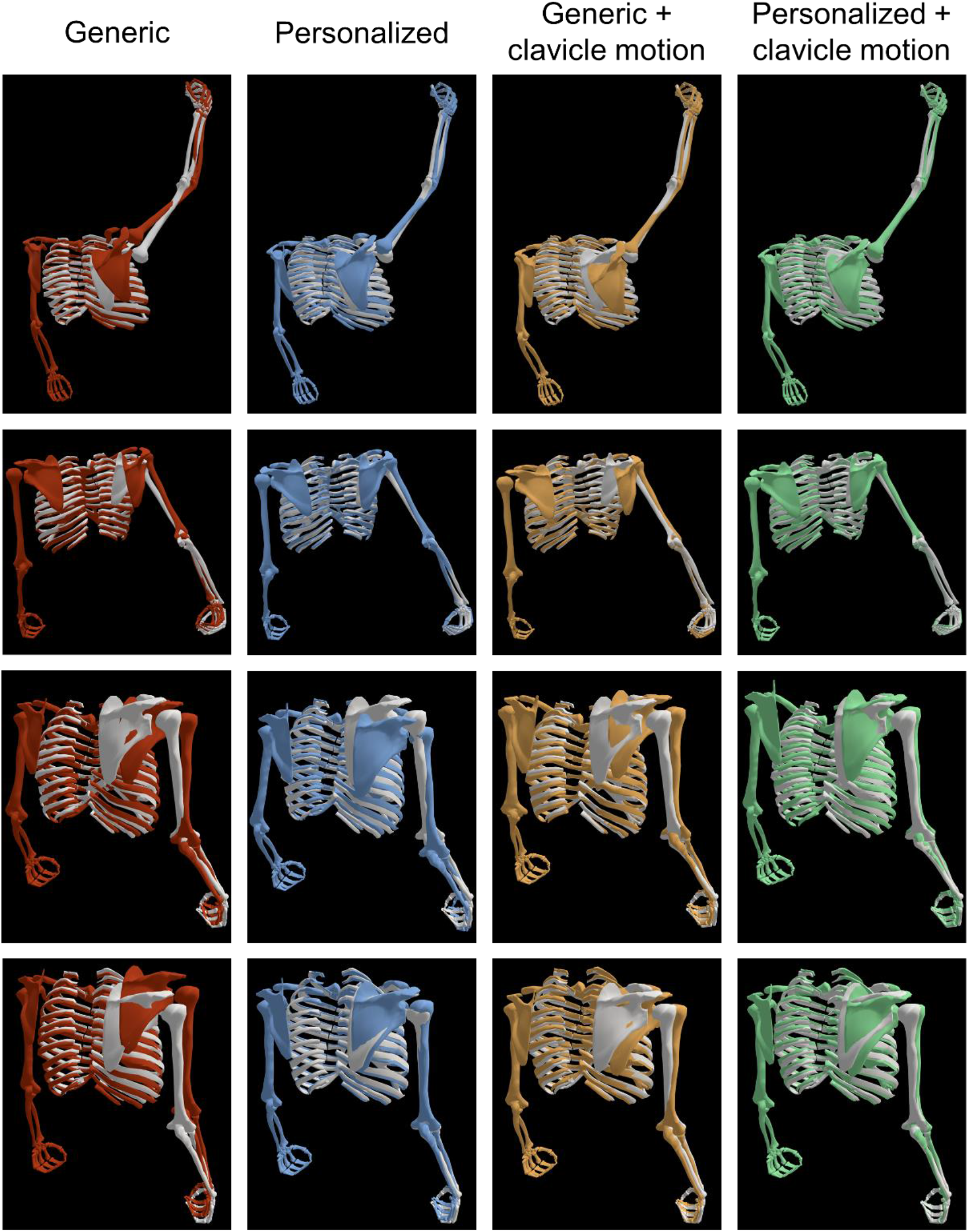
Models in different static poses. The white model represents the “target model” from computing inverse kinematics using skin markers and the palpated scapula markers. From left to right, the simulation results of the fully constrained generic model (red), the fully constrained model with personalized constraints (blue), the model with generic constraints and independent clavicle elevation motion (yellow), and the model with personalized constraints and independent clavicle elevation motion (green) are overlaid.

Quantitatively, both personalizing the shoulder constraints and allowing independent clavicle elevation reduced the average marker tracking error of all upper body markers including the palpated scapula markers (Fig. 4, *p* < 0.001 for both). Adding both modifications significantly reduced the marker RMS error compared to applying only model personalization (*t*(9) = −8.7, *p* < 0.001) or only independent clavicle elevation motion (*t*(9) = −4.1, *p* = 0.003). The personalized models achieved a significantly lower RMS error compared to the models with generic constraints in both scapula marker error (*p* < 0.001) and scapula axis-angle error (*p* < 0.001). Allowing independent clavicle elevation motion did not reduce the marker error (*p* = 0.278) or the axis-angle error (*p* = 0.476) in the scapula. We detected no significant interaction effect for the marker error (*p* = 0.101) or the axis-angle error (*p* = 0.522).

**Figure 4.**
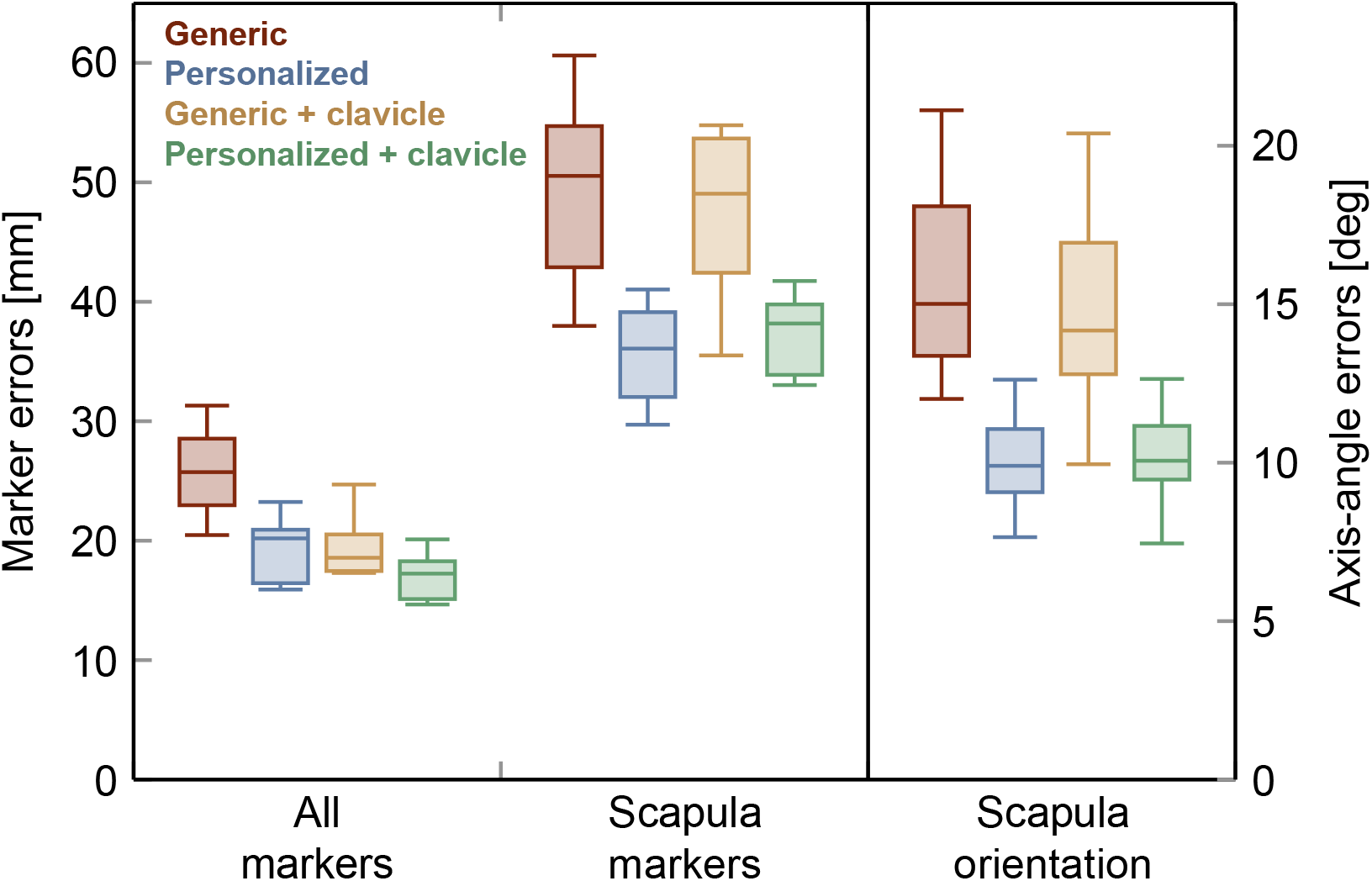
Static analysis. The box plots show the median over all subjects of the average errors of the simulated model markers compared with the 3D measurements of anatomical upper body markers and palpated scapula markers. The leftmost group of box plots shows the median of the root mean squared (RMS) error over all static trials of all subjects of all upper body markers including the scapula markers; the middle group shows the median of the RMS error for the scapula markers only; the rightmost group shows the median of the axis-angle orientation error of the scapula.

We observed similar quantitative results for the dynamic trials (Fig. 5). When considering all measured anatomical markers, both the constraint personalization and the independent clavicle elevation motion led to a significant reduction of the marker error (*p* < 0.001 for both). Applying both modifications to the model significantly reduced the marker RMS error of all markers compared to model personalization only (*t*(9) = −7.6, *p* < 0.001) or independent clavicle elevation motion only (*t*(9) = −3.0, *p* = 0.014). Scapula marker error and scapula axis-angle error were significantly reduced for the models with personalized constraints (*p* < 0.001 for both), but not for the models with independent clavicle elevation motion (*p* = 0.272 and *p* = 0.172). We observed a significant interaction effect for the scapula marker error (*p* = 0.048) but not for the axis-angle errors (*p* = 0.304). Applying both modifications to the model significantly increased the scapula marker RMS error compared to applying model personalization only (*t*(9) = 3.0, *p* = 0.015), and significantly decreased the scapula marker RMS error compared to independent clavicle elevation motion only (*t*(9) = −5.6, *p* < 0.001). We observed similar characteristics regarding the scapula, shoulder, and arm matching as for the static trials when visually comparing the models during a dynamic abduction trial at the best matching time points with the static target models (Fig. 6).

**Figure 5.**
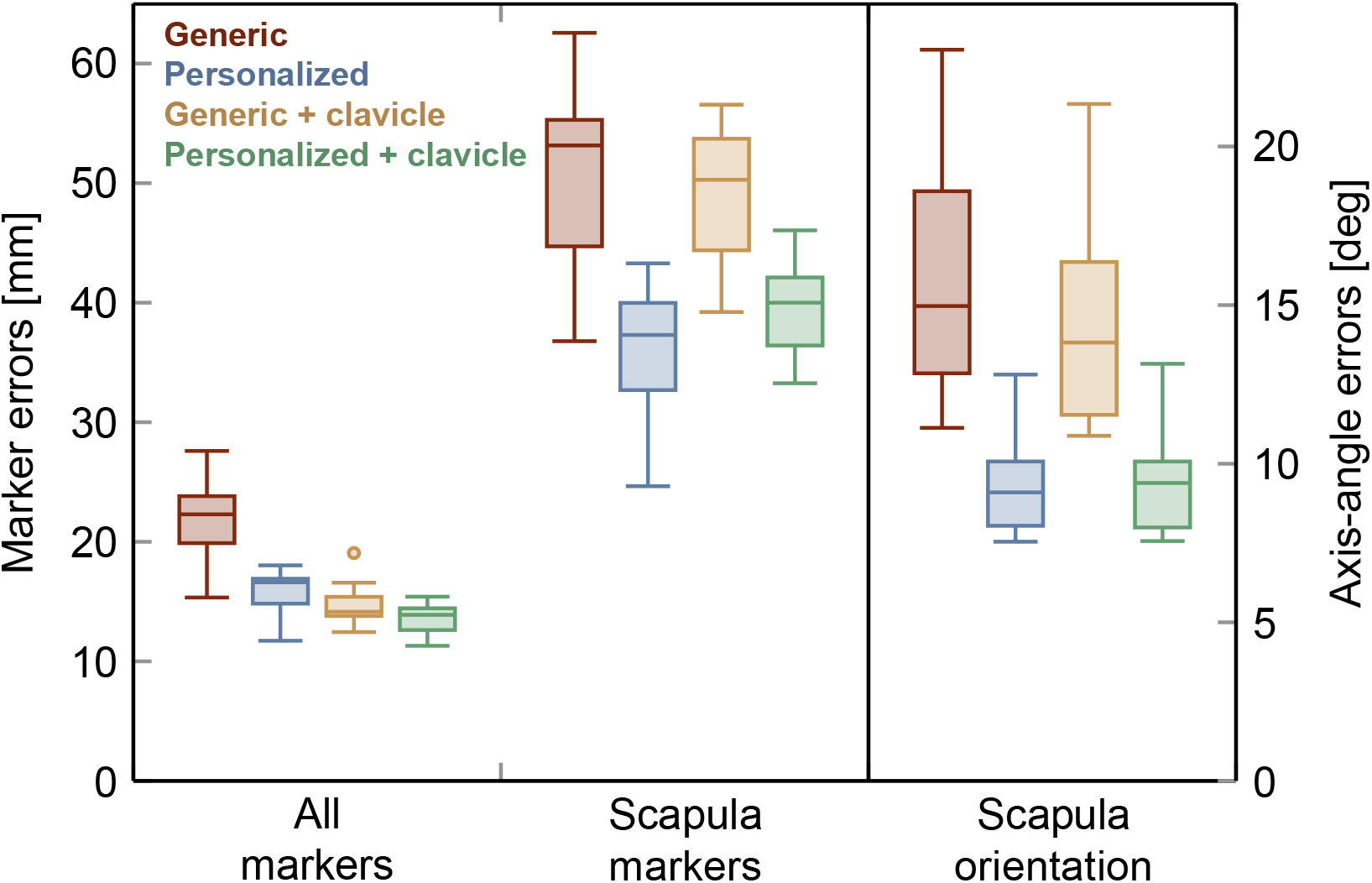
Dynamic analysis. The leftmost group of box plots show the median over all subjects of the root mean squared (RMS) error of the simulated model markers compared with the ground truth 3D measurements of anatomical upper body markers; the middle and rightmost group show the median of the scapula RMS error and the median of the axis-angle errors when comparing the scapula positions with the palpated scapula markers at the time points that best match the respective static trials.

**Figure 6.**
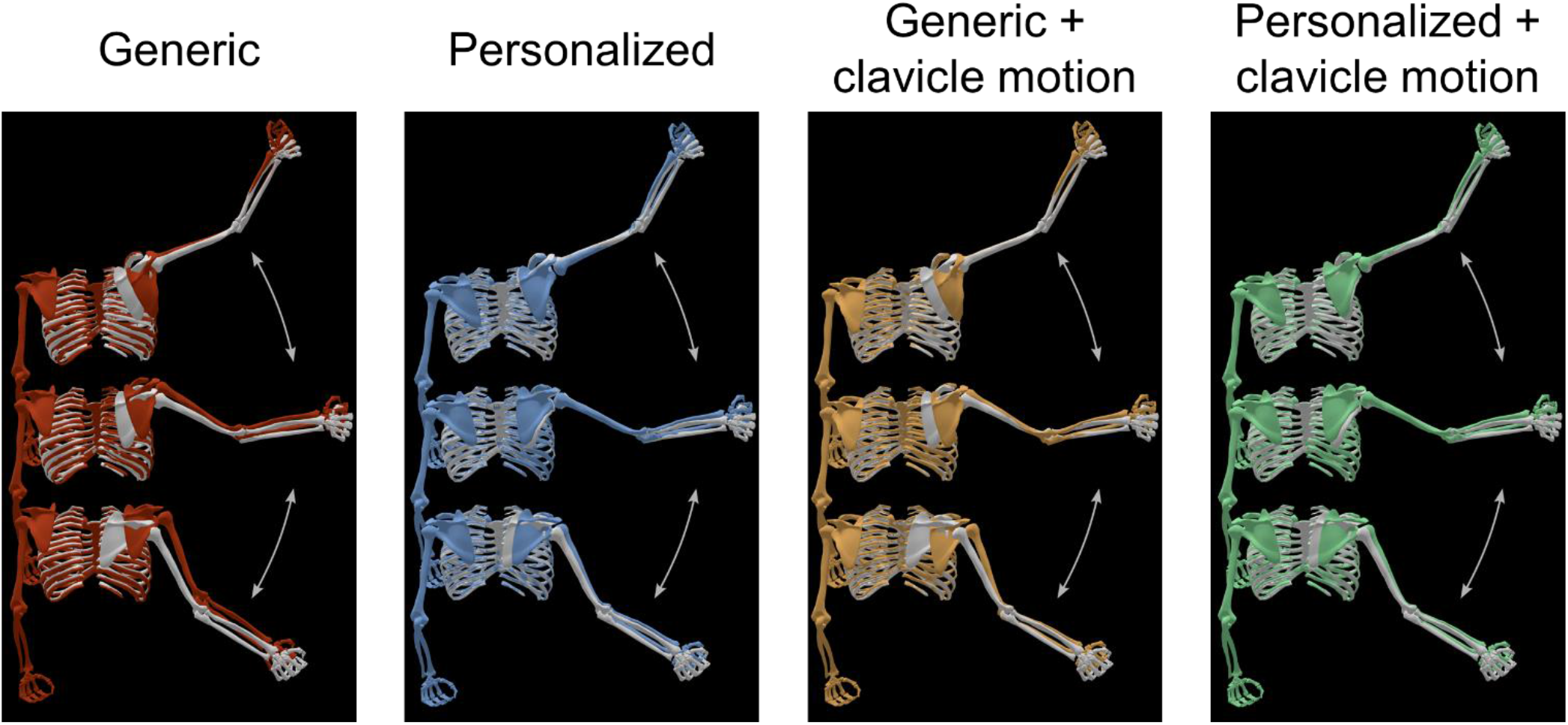
Models during dynamic abduction trial of one subject. The white model represents the best possible fit to the measured and palpated marker positions of the static trials at 45°, 90°, and 135° abduction, from bottom to top. The overlaid colored models represent a snapshot of the dynamic trial results at the time points that best match with the respective static trials. Generic model (red), personalized model (blue), generic model with independent clavicle elevation motion (yellow), and personalized model with independent clavicle elevation motion (green), are shown from left to right.

## 4. Discussion

Our results show that personalizing the shoulder rhythm of a computational upper body model reduces the errors in tracking scapular kinematics. We found that personalizing the shoulder rhythm constraints and allowing independent clavicle elevation motion improves the overall upper body marker tracking. However, the generic model with independent clavicle elevation only improves arm kinematic tracking and not scapular kinematic tracking. Applying both the constraint personalization and the independent clavicle motion can further reduce the overall upper body marker error while accurately tracking scapular kinematics.

It is infeasible to obtain measurements of the scapula orientation using palpation during dynamic motion. Therefore, we use palpated scapula orientations of the best matching static poses as an estimate of this measurement. However, we observe a slight deviation between the static and dynamic trials since it is not always possible to find an exact matching pose (Fig. 6). This is not surprising, as the participants performed, for example, the dynamic abduction trials with a slightly different shoulder flexion than the controlled static trials.

Options to obtain a ground truth measurement of scapular orientation during motion include bone pin and bi-plane fluoroscopy measurements (Giphart et al., 2013; Ivester et al., 2015; Ludewig et al., 2009). Using bone pins, scapula kinematics can be calculated by solving an optimization to minimize the differences between the measured and calculated scapula positions. Yanagawa et al. (2008) reported average errors of less than 2° for scapula upward rotation and less than 9° for the other two rotations, corresponding to an axis-angle error of 13°, which is comparable to our approach. A constrained computational model that represents the scapulothoracic articulation as a point sliding on an ellipsoid surface achieved an average upper body bone pin marker tracking error of below 10 mm (Seth et al., 2019); the scapula marker error and scapula axis-angle error computed with these authors’ published materials result in averages of 12 ± 4 mm and 9° ± 2°. Our method shows larger deviations in scapula position but estimates orientation similarly and does not require position measurements. Bi-plane fluoroscopy-based scapula orientation estimates achieve low errors of on average 3-5° per axis, corresponding to an axis-angle error of 5-8° (Nicholson et al., 2017a). As this technique is limited to a small scanning volume it is not practical for high range-of-motion activities.

A prior study has shown the benefits of individualizing shoulder rhythm dependencies by fitting linear models on 11 calibration poses to obtain the scapula orientation as a function of all glenohumeral angles and clavicle protraction and elevation angles (Nicholson et al., 2017b). They compute unconstrained inverse kinematics from marker positions directly and achieve average scapula angle errors of 6-7° per axis when comparing with fluoroscopy-based orientations, corresponding to an axis-angle error of 12°. Our approach uses a constrained model representing the underlying skeletal geometry in a multibody dynamics analysis (Hicks et al., 2015). This makes our inverse kinematics computations more robust to noise and ensures that nonphysiological motions like joint separation are prevented (Lu et al., 1999).

Fully constrained shoulder models face problems when the shoulder moves independently of the arms in elevation and depression, which we address by allowing independent clavicle elevation (bottom two rows of Fig. 3). Interestingly, in dynamic trials, adding independent clavicle elevation motion to the model with personalized constraints significantly increases the marker error compared to the fully constrained model. By independently elevating the clavicle, the model is able to track the arm markers and the acromioclavicular marker better, irrespective of scapular kinematics. This upward shift can increase the scapula marker errors. The estimated marker at the acromioclavicular joint is another potential contributor to the increase in scapula marker error. Nevertheless, we expect the independent clavicle elevation motion to become relevant during the movements of overhead athletes who intentionally perform elevation and depression.

This study sets the stage to analyze the kinematics and the kinetics of overhead motions. As many muscles attach to the scapula, the scapula orientation and position play a crucial role in understanding muscle lengths and moment arms. A realistic representation of scapular kinematics is critical to enable an accurate analysis of activations and forces of the muscles in the shoulder. Many studies still use a generic model as there is no practical and available way to apply personalization (Barnamehei et al., 2021; Hybois et al., 2019). We enable others to perform such personalized analyses by making our experimental data, OpenSim model, and methods publicly available (https://simtk.org/projects/shoulder-perso and https://github.com/stanfordnmbl/shoulder-personalization).

## Acknowledgements

This study was funded by the Joe and Clara Tsai Foundation through the Wu Tsai Human Performance Alliance.

## References

Al Borno, M., O’Day, J., Ibarra, V., Dunne, J., Seth, A., Habib, A., Ong, C., Hicks, J., Uhlrich, S., Delp, S., 2022. Opensense: An open-source toolbox for inertial-measurement-unit-based measurement of lower extremity kinematics over long durations. Journal of NeuroEngineering and Rehabilitation 19, 1–11.

Andersson, J.A., Gillis, J., Horn, G., Rawlings, J.B., Diehl, M., 2019. Casadi: a software framework for nonlinear optimization and optimal control. Mathematical Programming Computation 11, 1–36.

Baena-Raya, A., Soriano-Maldonado, A., Rodríguez-Pérez, M.A., GarcíaDe-Alcaraz, A., Ortega-Becerra, M., Jiménez-Reyes, P., García-Ramos, A., 2021. The force-velocity profile as determinant of spike and serve ball speed in top-level male volleyball players. PLOS ONE 16, e0249612.

Barnamehei, H., Ghomsheh, F.T., Cherati, A.S., Pouladian, M., 2021. Kinematic models evaluation of shoulder complex during the badminton overhead forehand smash task in various speed. Informatics in Medicine Unlocked 26, 100697.

Benjamini, Y., Hochberg, Y., 1995. Controlling the false discovery rate: A practical and powerful approach to multiple testing. Journal of the Royal Statistical Society: Series B (Methodological) 57, 289–300.

BenSaïda, A., 2014. Shapiro-wilk and shapiro-francia normality tests. https://www.mathworks.com/matlabcentral/fileexchange/13964-shapiro-wilk-and-shapiro-francia-normality-tests. [Online; Retrieved January 10, 2024].

Betts, J.T., 2010. Practical Methods for Optimal Control and Estimation Using Nonlinear Programming, Second Edition. Society for Industrial and Applied Mathematics.

De Groot, J.H., Brand, R., 2001. A three-dimensional regression model of the shoulder rhythm. Clinical Biomechanics 16, 735–743.

Delp, S.L., Anderson, F.C., Arnold, A.S., Loan, P., Habib, A., John, C.T., Guendelman, E., Thelen, D.G., 2007. Opensim: open-source software to create and analyze dynamic simulations of movement. IEEE transactions on biomedical engineering 54, 1940–1950.

Dembia, C.L., Bianco, N.A., Falisse, A., Hicks, J.L., Delp, S.L., 2020. Opensim moco: Musculoskeletal optimal control. PLOS Computational Biology 16, e1008493.

Doorenbosch, C.A.M., Harlaar, J., DirkJan, V., 2003. The globe system: An unambiguous description of shoulder positions in daily life movements. Journal of Rehabilitation Research and Development 40, 147–156.

Giphart, J.E., Brunkhorst, J.P., Horn, N.H., Shelburne, K.B., Torry, M.R., Millett, P.J., 2013. Effect of plane of arm elevation on glenohumeral kinematics: A normative biplane fluoroscopy study. Journal of Bone and Joint Surgery 95, 238–245.

Grewal, T.J., Dickerson, C.R., 2013. A novel three-dimensional shoulder rhythm definition that includes overhead and axially rotated humeral postures. Journal of Biomechanics 46, 608–611.

Hashimoto, Y., Nagami, T., Yoshitake, S., Nakata, H., 2023. The relationship between pitching parameters and release points of different pitch types in major league baseball players. Frontiers in Sports and Active Living 5, 1113069.

Hicks, J.L., Uchida, T.K., Seth, A., Rajagopal, A., Delp, S.L., 2015. Is my model good enough? best practices for verification and validation of musculoskeletal models and simulations of movement. Journal of Biomechanical Engineering 137, 020905.

Holzbaur, K.R., Murray, W.M., Delp, S.L., 2005. A model of the upper extremity for simulating musculoskeletal surgery and analyzing neuromuscular control. Annals of Biomedical Engineering 33, 829–840.

Hybois, S., Puchaud, P., Bourgain, M., Lombart, A., Bascou, J., Lavaste, F., Fodé, P., Pillet, H., Sauret, C., 2019. Comparison of shoulder kinematic chain models and their influence on kinematics and kinetics in the study of manual wheelchair propulsion. Medical Engineering & Physics 69, 153–160.

Ivester, J.C., Cyr, A.J., Harris, M.D., Kulis, M.J., Rullkoetter, P.J., Shelburne, K.B., 2015. A reconfigurable high-speed stereo-radiography system for sub-millimeter measurement of in vivo joint kinematics. Journal of Medical Devices, Transactions of the ASME 9.

Lu, T.W., O’Connor, J.J., 1999. Bone position estimation from skin marker co-ordinates using global optimisation with joint constraints. Journal of Biomechanics 32, 129–134.

Ludewig, P.M., Phadke, V., Braman, J.P., Hassett, D.R., Cieminski, C.J., Laprade, R.F., 2009. Motion of the shoulder complex during multiplanar humeral elevation. Journal of Bone and Joint Surgery 91, 378–389.

Makhsous, M., 1999. Improvements, validation and adaptation of a shoulder model. Goteborg: Chalmers University of Technology, 1–43.

Monnet, T., Desailly, E., Begon, M., Vallee, C., Lacouture, P., 2007. Comparison of the score and ha methods for locating in vivo the glenohumeral joint centre. Journal of Biomechanics 40, 3487–3492.

Nicholson, K.F., Richardson, R.T., Miller, F., Richards, J.G., 2017a. Determining 3d scapular orientation with scapula models and biplane 2d images. Medical Engineering and Physics 41, 103–108.

Nicholson, K.F., Richardson, R.T., Rapp, E.A., Quinton, R.G., Anzilotti, K.F., Richards, J.G., 2017b. Validation of a mathematical approach to estimate dynamic scapular orientation. Journal of Biomechanics 54, 101–105.

Pascoal, A.G., Ribeiro, A., Infante, J., 2023. Scapular resting posture and scapulohumeral rhythm adaptations in volleyball players: Implications for clinical shoulder assessment in athletes. MDPI Sports, Vol. 11, 114.

Ribeiro, A., Pascoal, A.G., 2013. Resting scapular posture in healthy overhead throwing athletes. Manual Therapy 18, 547–550.

Saul, K.R., Hu, X., Goehler, C.M., Vidt, M.E., Daly, M., Velisar, A., Murray, W.M., 2015. Benchmarking of dynamic simulation predictions in two software platforms using an upper limb musculoskeletal model. Computer methods in biomechanics and biomedical engineering 18, 1445.

Seabold, S., Perktold, J., 2010. statsmodels: Econometric and statistical modeling with python, in: 9th Python in Science Conference, pp. 92–96.

Seth, A., Dong, M., Matias, R., Delp, S., 2019. Muscle contributions to upper-extremity movement and work from a musculoskeletal model of the human shoulder. Frontiers in Neurorobotics 13.

Seth, A., Hicks, J.L., Uchida, T.K., Habib, A., Dembia, C.L., Dunne, J.J., Ong, C.F., DeMers, M.S., Rajagopal, A., Millard, M., et al., 2018. Opensim: Simulating musculoskeletal dynamics and neuromuscular control to study human and animal movement. PLOS Computational Biology 14, e1006223.

Seth, A., Matias, R., Veloso, A.P., Delp, S.L., 2016. A biomechanical model of the scapulothoracic joint to accurately capture scapular kinematics during shoulder movements. PLOS ONE 11, e0141028.

Van Der Helm, F.C.T., 1994. A finite element musculoskeletal model of the shoulder mechanism. Journal of Biomechanics 21, 551–569.

Virtanen, P., Gommers, R., Oliphant, T.E., Haberland, M., Reddy, T., Cournapeau, D., Burovski, E., Peterson, P., Weckesser, W., Bright, J., van der Walt, S.J., Brett, M., Wilson, J., Millman, K.J., Mayorov, N., Nelson, A.R.J., Jones, E., Kern, R., Larson, E., Carey, C.J., Polat, I., Feng, Y., Moore, E.W., VanderPlas, J., Laxalde, D., Perktold, J., Cimrman, R., Henriksen, I., Quintero, E.A., Harris, C.R., Archibald, A.M., Ribeiro, A.H., Pedregosa, F., van Mulbregt, P., SciPy 1.0 Contributors, 2020. SciPy 1.0: Fundamental Algorithms for Scientific Computing in Python. Nature Methods 17, 261–272.

Wächter, A., Biegler, L.T., 2006. On the implementation of an interior-point filter line-search algorithm for large-scale nonlinear programming. Mathematical programming 106, 25–57.

Wu, G., Van Der Helm, F.C., Veeger, H.E., Makhsous, M., Roy, P.V., Anglin, C., Nagels, J., Karduna, A.R., McQuade, K., Wang, X., Werner, F.W., Buchholz, B., 2005. Isb recommendation on definitions of joint coordinate systems of various joints for the reporting of human joint motion - part ii: Shoulder, elbow, wrist and hand. Journal of Biomechanics 38, 981–992.

Xu, X., Dickerson, C.R., hua Lin, J., McGorry, R.W., 2016. Evaluation of regression-based 3-d shoulder rhythms. Journal of Electromyography and Kinesiology 29, 28–33.

Xu, X., hua Lin, J., McGorry, R.W., 2014. A regression-based 3-d shoulder rhythm. Journal of Biomechanics 47, 1206–1210.

Xu, X., Lin, J.H., McGorry, R.W., 2012. Coordinate transformation between shoulder kinematic descriptions in the holzbaur et al. model and isb sequence. Journal of Biomechanics 45, 2715–2718.

Yanagawa, T., Goodwin, C.J., Shelburne, K.B., Giphart, J.E., Torry, M.R., Pandy, M.G., 2008. Contributions of the individual muscles of the shoulder to glenohumeral joint stability during abduction. Journal of Biomechanical Engineering 130(2), 021024.

