## Appendix for "Personalizing a computational upper body model improves kinematic tracking in high range-of-motion arm movements"

### Appendix A. Coordinate system definitions

Table A.1: Coordinate system definitions of the generic model (De Groot and Brand, 2001) and the coordinate system definitions recommended by the International Society of Biomechanics (Wu et al., 2005).

| **Body/Joint** | **De Groot and Brand (2001)** | **ISB recommendations (Wu et al., 2005)** |
|---|---|---|
| Global | $\boldsymbol{x}_G$: Horizontally to the right<br>$\boldsymbol{y}_G$: Vertically upward<br>$\boldsymbol{z}_G$: Horizontally backward | Not defined |
| Thorax | $\boldsymbol{o}_T$: *IJ*<br>$\boldsymbol{x}_T$: ⊥ on plane [*IJ, PX, C7*]<br>$\boldsymbol{y}_T$: *IJ - PX*<br>$\boldsymbol{z}_T$: ⊥ on plane $(\boldsymbol{x}_T, \boldsymbol{y}_T)$ | $\boldsymbol{o}_T$: *IJ*<br>$\boldsymbol{y}_T$: 0.5(*IJ* + *C7*) – 0.5(*PX* + *T8*)<br>$\boldsymbol{z}_T$: ⊥ on plane [*IJ, C7,* 0.5(*PX* + *T8*)]<br>$\boldsymbol{x}_T$: ⊥ on plane $(\boldsymbol{z}_T, \boldsymbol{y}_T)$ |
| Clavicle | $\boldsymbol{o}_C$: *SC*<br>$\boldsymbol{x}_C$: (*AC – SC*)<br>$\boldsymbol{z}_C$: ⊥ on plane $(\boldsymbol{x}_C, \boldsymbol{y}_G)$<br>$\boldsymbol{y}_C$: ⊥ on plane $(\boldsymbol{x}_C, \boldsymbol{z}_C)$ | $\boldsymbol{o}_C$: *SC*<br>$\boldsymbol{z}_C$: (*AC – SC*)<br>$\boldsymbol{x}_C$: ⊥ on plane $(\boldsymbol{z}_C, \boldsymbol{y}_T)$<br>$\boldsymbol{y}_C$: ⊥ on plane $(\boldsymbol{x}_C, \boldsymbol{z}_C)$ |
| Scapula | $\boldsymbol{o}_S$: *AC*<br>$\boldsymbol{x}_S$: (*AA – TS*)<br>$\boldsymbol{z}_S$: ⊥ on plane [*AA, TS, AI*]<br>$\boldsymbol{y}_S$: ⊥ on plane $(\boldsymbol{x}_S, \boldsymbol{z}_S)$ | $\boldsymbol{o}_S$: *AC*<br>$\boldsymbol{x}_S$: ⊥ on plane [*AA, TS, AI*]<br>$\boldsymbol{z}_S$: (*AA – TS*)<br>$\boldsymbol{y}_S$: ⊥ on plane $(\boldsymbol{x}_S, \boldsymbol{z}_S)$ |
| Humerus | $\boldsymbol{o}_H$: *GH*<br>$\boldsymbol{y}_H$: (*GH – EP*) with *EP* = 0.5(*EM* + *EL*)<br>$\boldsymbol{z}_H$: ⊥ on plane [*GH, EM, EL*]<br>$\boldsymbol{x}_H$: ⊥ on plane $(\boldsymbol{y}_H, \boldsymbol{z}_H)$ | $\boldsymbol{o}_H$: *GH*<br>$\boldsymbol{y}_H$: (*GH – EP*) with *EP* = 0.5(*EM* + *EL*)<br>$\boldsymbol{x}_H$: ⊥ on plane [*GH, EM, EL*]<br>$\boldsymbol{z}_H$: ⊥ on plane $(\boldsymbol{y}_H, \boldsymbol{x}_H)$ |
| Clavicle<br>relative to thorax | Same as clavicle | $\boldsymbol{o}_{SC}$: *SC*<br>$\boldsymbol{x}_{SC}$: $\boldsymbol{y}_T$<br>$\boldsymbol{z}_{SC}$: $\boldsymbol{z}_C$<br>$\boldsymbol{y}_{SC}$: ⊥ on plane $(\boldsymbol{x}_{SC}, \boldsymbol{z}_{SC})$ |
| Scapula<br>relative to thorax | Same as scapula | $\boldsymbol{o}_{AC}$: *AC*<br>$\boldsymbol{x}_{AC}$: $\boldsymbol{y}_T$<br>$\boldsymbol{z}_{AC}$: $\boldsymbol{z}_S$<br>$\boldsymbol{y}_{AC}$: ⊥ on plane $(\boldsymbol{x}_{AC}, \boldsymbol{z}_{AC})$ |
| Humerus<br>relative to thorax | Same as humerus | $\boldsymbol{o}_{GH}$: *GH*<br>$\boldsymbol{x}_{GH}$: $\boldsymbol{y}_T$<br>$\boldsymbol{z}_{GH}$: $\boldsymbol{y}_H$<br>$\boldsymbol{y}_{GH}$: $\boldsymbol{x}_H$ |

IJ: Incisura jugularis, PX: Xiphoid process, C7: Seventh cervical vertebra, T8: Eigth thoracic vertebra, SC: Sternoclavicular joint center, AC: Acromioclavicular joint center, AA: Angulus acromialis, AI: Angulus inferior, TS: Trigonum spinae, GH: Glenohumeral joint center, EM: medial epicondyle, EL: lateral epicondyle.

#### Appendix B. Simulation details

We use OpenSim Moco (Dembia et al., 2020) to compute dynamically-constrained inverse kinematics by solving tracking optimization problems. In the following descriptions, OpenSim and Moco class names are denoted with italics. We use the *MocoTrack* tool to solve for both a motion and actuator controls that minimize the error compared to an observed motion in addition to control effort.

##### Models

The simulations were performed for the four tested models: the fully constrained generic model with the bivariate constraints, the fully constrained model with personalized constraints, the model with generic bivariate constraints and independent clavicle elevation, and the model with both personalized constraints and independent clavicle elevation. We added torque actuators to each coordinate of the model. The three actuators for the rotational coordinates between the ground and thorax had an optimal force of 100 *Nm* and the three actuators for the translational ground-thorax coordinates had optimal forces between 1000 *N* and 2000 *N*. We added activation coordinate actuators for all other 40 coordinates with an optimal force of 50 *Nm* and a 0.025 *s* activation time constant.

##### Cost function

The cost function consisted of two terms with individual weights that minimized control effort and tracked experimental markers (Eq. B.1). Each term is integrated over the duration of the trial with initial time $t_0$ and final time $t_f$. In the following equations, bold symbols represent vector quantities.

$$J = w_1 \, J_{\text{effort}} + w_2 \, J_{\text{track}} \tag{B.1}$$

We used *MocoControlGoal* to minimize the sum of squared torque actuator controls $e_i$, integrated over the trial duration (Eq. B.2).

$$J_{\text{effort}} = \int_{t_0}^{t_f} \sum_i^{46} e_i^2 \, dt \tag{B.2}$$

We used *MocoMarkerTrackingGoal* to minimize the sum of weighted squares between the 22 experimental markers, $\boldsymbol{x}_i^{\text{exp}}$, and model markers, $\boldsymbol{x}_i$ (Eq. B.3).

$$J_{\text{track}} = \int_{t_0}^{t_f} \sum_i^{22} \omega_i \left\| \boldsymbol{x}_i^{\text{exp}} - \boldsymbol{x}_i \right\|^2 dt \qquad \text{(B.3)}$$

The weights for the markers, $\omega_i$, were chosen such that torso markers were weighted highest to prevent excessive lumbar bending in case the shoulder definition did not allow for a good tracking of both arm and torso markers. When tracking the clavicle elevation, we weighted the virtual marker at the acromioclavicular joint higher than the markers on the arms to achieve smooth shoulder elevation changes. The tracking markers on the upper arms and forearms were weighted lowest because they were placed on soft tissue and not as reliably measured as anatomical markers. All individual marker weights are listed in Table B.2.

Table B.2: Marker weights per marker group.

| **Markers** | **Weights** |
|---|---|
| Torso markers | 10 |
| Elbow, wrist, and hand markers | 1 |
| Tracking markers (upper arm and forearm) | 0.1 |
| Markers on flat part of the acromion | 1 if clavicle elevation constrained<br>0 if clavicle elevation tracked |
| Virtual markers at acromioclavicular joints | 0 if clavicle elevation constrained<br>3 if clavicle elevation tracked |

The weights for each cost function term listed in Table B.3 were chosen manually with the goal of tracking the measured markers as closely as possible while enforcing the skeletal dynamics of the shoulder motion.

Table B.3: Cost function weights.

| **Weight** | **Value** |
|---|---|
| $w_1$ | 0.01 |
| $w_2$ | 100 |

#### Problem constraints

Since we knew that the participants performed slow controlled movements and did not move their feet throughout the trials, we implemented constraints on the coordinate speeds (Table B.4).

Table B.4: Speed constraints for the joint coordinates.

| **Joint** | **Speed range [min, max]** |
|---|---|
| ground thorax | [-0.5, 0.5] |
| sternoclavicular | [-1, 1] |
| acromioclavicular | [-1, 1] |
| glenohumeral | [-2, 2] |
| elbow | [-2, 2] |
| radioulnar | [-2, 2] |
| radiocarpal | [-2, 2] |

We used an initial guess for the first frame of each processed trial from an inverse kinematics (IK) computation using the marker weights listed in Table B.2. All joint coordinates were constrained to lie within the range of ± 0.05 *rad* of the values in the IK solution for rotational coordinates and ± 0.05 *m* of the values in the IK solution for translational coordinates. If the IK solution value ± 0.05 resulted in a value lower than the minimum allowed value or higher than the maximal allowed value for a certain coordinate, the range was capped at that minimal or maximal allowed value.

#### Solver settings

We solved each problem in Moco using *MocoCasADiSolver* which uses CasADi to transcribe each optimal control problem into a non-linear program (NLP) (Andersson et al., 2019). Each NLP is then solved by IPOPT (Wächter and Biegler, 2006). We used a Hermite-Simpson collocation scheme with mesh intervals at every 100 milliseconds. We determined the discretization for the simulations based on a sensitivity analysis using three representative dynamic trials from our dataset. We compared objective values and joint angle trajectories from simulations generated with mesh intervals at every 50, 100, 200, and 300 milliseconds. While results did not vary greatly between the tested discretizations, we selected a discretization with mesh intervals every 100 milliseconds as a compromise between dynamic accuracy and computational speed in each simulation, which were atypically long for dynamic optimization (often greater than 10 seconds). Each problem was solved using a constraint tolerance of $1e{-}2$ and a convergence tolerance of $1e{-}2$. We used a forward difference scheme to compute function derivatives in CasADi since this reduced optimization times but did not negatively affect problem convergence. A list of important solver settings can be found in Table B.5.

Table B.5: CasADi solver settings.

| **Setting** | **Value** |
|---|---|
| `transcription_scheme` | `‘hermite-simpson’` |
| `optim_constraint_tolerance` | `1e-2` |
| `optim_convergence_tolerance` | `1e-2` |
| `num_mesh_intervals` | `t/0.1 s` |
| `multibody_dynamics_mode` | `‘explicit’` |
| `optim_finite_difference_scheme` | `‘forward’` |
| `scale_cariables_using_bounds` | `false` |
